# Studying Pathogens Degrades BLAST-based Pathogen Identification

**DOI:** 10.1101/2022.07.12.499705

**Authors:** Jacob Beal, Adam Clore, Jeff Manthey

**Author notes:** Contributing authors.

## Abstract

As synthetic biology becomes increasingly capable and accessible, it is likewise increasingly critical to be able to make accurate biosecurity determinations regarding the pathogenicity or toxicity of particular nucleic acid or amino acid sequences. At present, this is typically done using the BLAST algorithm to determine the best match with sequences in the NCBI databases. Neither BLAST nor the NCBI databases, however, are actually designed for biosafety determination. Critically, taxonomic errors or ambiguities in the NCBI databases can also cause errors in BLAST-based taxonomic categorization. With heavily studied taxa and frequently used biotechnology tools, even low frequency taxonomic categorization issues can lead to high rates of errors in biosecurity decision-making. Here we focus on the implications for false positives, finding that NCBI BLAST will now incorrectly categorize a number of commonly used biotechnology tool sequences as the pathogens or toxins with which they have been used. Paradoxically, this implies that problems are expected to be most acute for the pathogens and toxins of highest interest and the most widely used biotechnology tools. We thus conclude that biosecurity tools should shift away from BLAST against NCBI and towards new methods that are specifically tailored for biosafety purposes.

## 1 Introduction

Advancements in synthetic biology are continuing to rapidly increase both organism engineering capabilities and the accessibility of those capabilities to a broad range of potential actors [1]. This necessarily increases the potential biosecurity risk from misuse of these capabilities, such as deliberate or accidental use of gene synthesis and gene editing to produce dangerous pathogenic organisms or toxins [2–6]. Moreover, the gene and protein sequences for dangerous organisms and toxins are readily available, e.g., in the NCBI databases and similar public scientific resources.

As a consequence, it is becoming increasingly critical to be able to make rapid and accurate determinations of the risk posed by particular nucleic acid sequences or amino acid sequences. Organizations like the International Gene Synthesis Consortium (IGSC) and its members need to make these determinations in order to determine whether or not the materials a customer has ordered are subject to legal controls and whether or not to fill that order [7, 8]. Similar concerns are present for organizations that design or edit organisms, for DNA depositories such as AddGene or the iGEM registry, and many other organizations as well.

At present, the most common means of evaluating the risk posed by a nucleic or amino acid sequence is to use the BLAST algorithm suite [9] to determine which sequences in the NCBI databases^1^ are most closely related to the sequence under consideration. Typically, if the sequence is found to be more closely related to some controlled pathogen or toxin than to any non-controlled sequence, then the sequence is considered controlled, unless an expert judges the sequence to fall into an exception category such as being a common “housekeeping gene” [2].

Neither BLAST nor the NCBI databases, however, are specifically designed for the purpose of determining whether a sequence is from dangerous pathogen or toxin. Taxonomic classification ambiguities that are allowed in the NCBI databases to support their intended purposes can result in misclassification, as illustrated in Figure 1. For example, if a tag is added to a protein in order to enable a crystallization study, then the modified protein will generally be appropriately categorized under the original taxon, as that is the subject of the study. Likewise, a virus modified to include a fluorescent reporter might still be categorized under the original virus. Various forms of horizontal transfer can create similar chimerism naturally as well. These categorizations are not necessarily incorrect, as the sequence is indeed generally most relevant to the taxon to which it has been assigned.

**Fig. 1.**
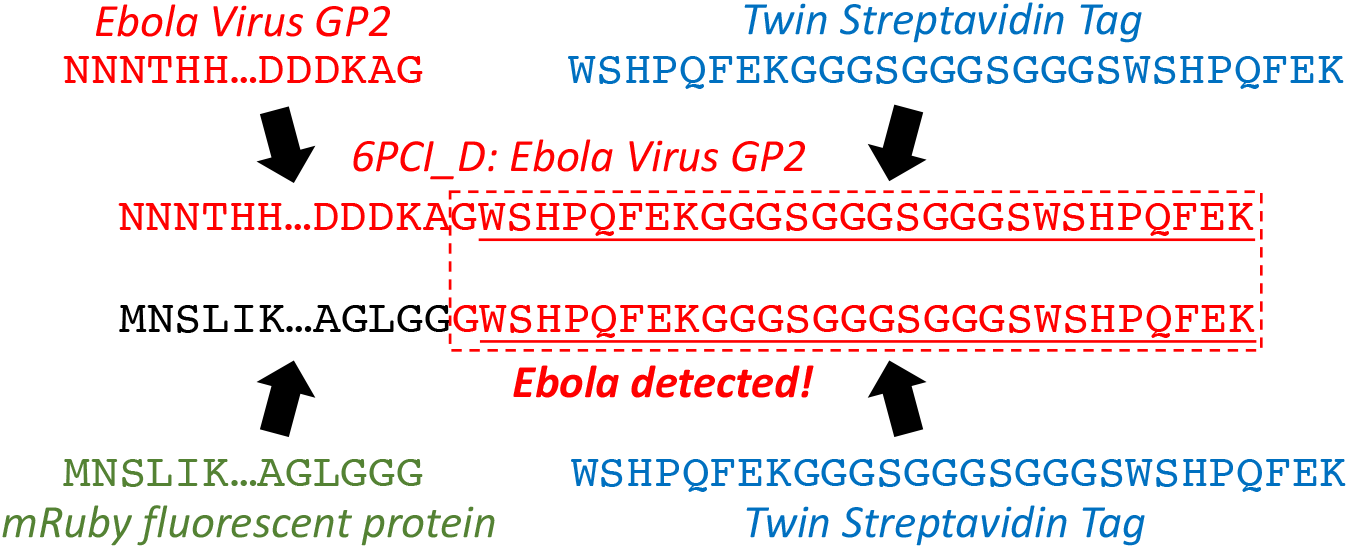
NCBI databases allow categorization of chimeric material under the taxa most relevant to the study. For example, NCBI protein accession 6PCI_D is classified as Ebola virus since it is the Ebola virus GP2 protein, studied with the aid of an appended twin streptavidin tag. BLAST matching of new material can then misidentify its taxa by matching against the chimeric material. For example, when the twin streptavidin tag is added to the mRuby protein, BLAST on its 3’ end produces a best match with 6PCI_D, since they share their last amino acid before the tag, thus identifying the sequence as controlled Ebola virus material, despite it being completely unrelated.

Assigning a chimeric sequence to a non-chimeric taxon, however, necessarily means that the assigned taxon now includes some amount of sequence materials derived from other taxa. These chimeric materials can then be matched by BLAST, identifying both the original taxa and the chimeric taxa. Depending on the specifics of the chimerism and the taxa involved, this can either cause benign material to appear dangerous (as in the example in Figure 1) or cause dangerous material to appear benign. For common subjects of study, such as model organisms or important pathogens and toxins, the relative number of sequences assigned to the taxon may be quite high indeed (for example, as of this writing, SARS-CoV-2 sequences in NCBI comprise 59.5% of all viral nucleotide sequences and 75.3% of all viral protein sequences), and chimeric usages may come to outweigh the original or even to crowd it entirely out of BLAST results. Moreover, many biotechnology tools are originally categorized as artificial sequences, a polyphyletic category that by its nature cannot be used for determination of potential pathogen or toxin content.

In this case study, we investigate whether such taxonomic ambiguities can, in fact, result in incorrect determinations of sequence risk based on the results of BLAST against NCBI databases. For reasons of information safety, we specifically focus on false positives, in which benign biotechnology tools are mis-categorized as dangerous pathogens or toxins.

## 2 Results

For this case study, we selected seven protein sequences for common biotechnology tools, ranging in length from 18 to 231 amino acids. One of these is the T4 foldon, a short sequence from an *E. coli* bacteriphage that has a long history of use for stabilization of proteins (e.g. [10, 11]), including applications in vaccines (e.g., [12–14]). Three others are protein tags used for purification: a peptide signal for secretion [15] and the streptavidin tag [16] either coupled with a PreScission protease target or in a twin streptaptividin configuration [17]. The final three are reporter sequences, which have greater length: the first 60 amino acids of a GFP fluorescent protein, the NanoLuc luciferase [18], and the ZsGreen fluorescent protein [19]. Sequences are provided in Supplementary S1.

This case study thus coves multiple different common classes of proteinbased tool, with sequence sizes that extend from well above the current standard screening threshold of 200 nucleic acids base pairs [2, 7] down to just above the screening threshold of 50 base pairs recently proposed by the US DHHS [20].

To analyze the potential impact of chimeric material on determination of controlled pathogen/toxin status for these genetic tools, we first ran a protein BLAST for each of the seven amino acid sequences and examined the taxonomic categorization of the results (see Methods for details). Since each of these sequences is widely used, BLAST returns many sequences that are equal “best matches” with respect to maximum bit-score (the metric typically used for determination of control status), including some that match material from sequences categorized as controlled pathogens or toxins. The fraction of matches for each sequence to controlled material (Figure 2 center) spans a broad range, from 1.6% for NanoLuc to a remarkable 93% for the T4 foldon, with an overall average of 33.8%. Since the results for each sequence also contain non-controlled sequences with equal best-match scores, these sequences would all appropriately be assessed as non-controlled.

**Fig. 2.**
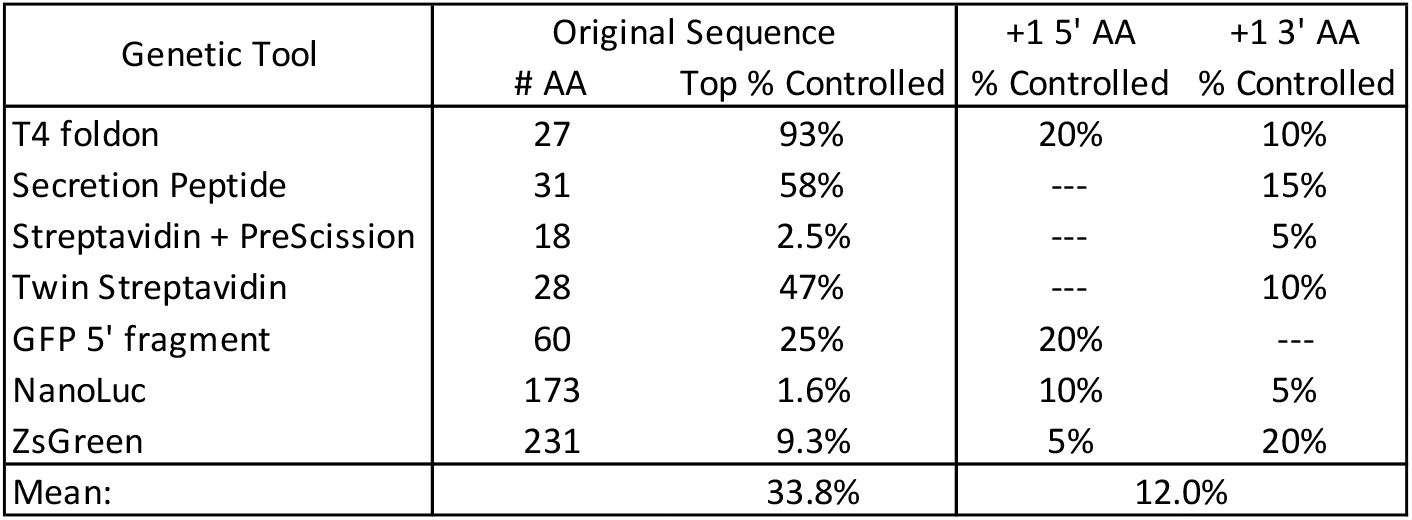
For each test sequence, BLAST analysis found many matches with the same maximum bit-score. Of these tied matches, the fraction of sequences classified as controlled pathogens or toxins varies widely. When the sequence was extended with each of the 20 possible amino acids added to its 3’ or 5’ end, between 5% and 20% (1-4 of the options) have best matchess consisting only of controlled pathogens or toxins. For some sequences, only one side was analyzed: the secretion and streptavidin tags are typically used at the 5’ end of a sequence, and the GFP 5’ fragment is constrained to follow the GFP sequence on its 3’ side.

The high volume of controlled sequence matches, however, indicates that classification is likely to be fragile. Even a single additional amino acid that matches better to a controlled sequence than a non-controlled sequence can change the classification, as in the case shown in Figure 1. To study this fragility, we ran BLAST against sequences extended by a single amino acid at the 5’ and/or 3’ end, systematically testing each of the 20 possible options for an additional canonical amino acid. The protein tags are typically added to the 5’ end, so only 3’ extensions were tested for these sequences, and the GFP fragment contains only the 3’ portion of the protein, so only 5’ extensions were tested for that sequence. As predicted, with the addition of a single additional amino acid, the previous ties for best match were indeed often broken in favor of controlled sequences (Figure 2 right), with an average of 12% of all extensions having a best match only to controlled sequences.

The taxonomic classification of each best match to a controlled sequence is provided in the “1 AA flanker” table in Supplementary 3. The most frequent match was to the SARS-CoV-2 virus, which was a best match for 11 out of the 24 sequences with controlled best matches. Other controlled taxa were mainly viral—high pathogenicity avian influenza, Newcastle disease virus, Vesicular stomatitis virus, Yellow fever virus, and Ebola virus—but one match was to the *Clostridium botulinum* hemagglutinin protein, part of the botulinum neurotoxin (BoNT) complex [21].

Since some of these sequences are much smaller than the commonly used 200 base pair threshold, one might wonder whether these results would still hold when considering the sequence in a larger context. To test this hypothesis, we selected one of the controlled sequences from each of the T4 foldon extensions and from the tag extensions, then generated for each selected sequence 10 extensions with random sequences of 60 additional amino acids and ran BLAST against these 50 extended sequences. The taxonomic classification of each best match to a controlled sequence is provided in the “Random Extension” table in Supplementary 3. In some cases, the random additional material did indeed break the best match relationship (Figure 3), though in two of these the new best match was a controlled sequence from a different taxa. For all except the secretion peptide, however, the single amino acid extension strongly predicted the classification of the larger sequence, with at least half of the random sequences still best matching to controlled sequences, and overall 27 out of the 50 extended sequences. In sum, then, these results indicate that any inclusion of a genetic tool in a sequence classified as a controlled pathogen or toxin results in a significant chance that other uses of the same tool will be classified as controlled as well.

**Fig. 3.**
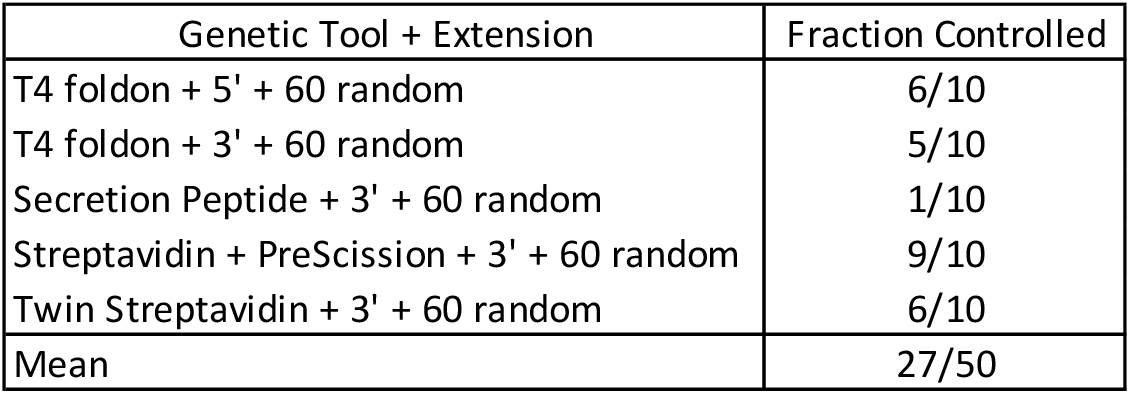
For each short sequence, a single amino-acid extension classified as pathogenic was further extended with 10 different 60 additional random animo acid sequences. For all except the secretion peptide, the single amino acid strongly predicted the classification of the extended sequence, with at least half of the random sequences also being classified as a controlled pathogen.

## 3 Discussion

These results demonstrate a serious problem in the use of BLAST against NCBI databases to identify pathogens. Although only a handful of sequences were selected for this case study, in our current biosecurity screening deployment we have seen these same issues commonly arise with a wide range of other common biotechnology tools, including antibodies, sequencing adapters, antibiotic selection markers, plasmid vectors, and promoters. A similar dynamic applies for the more dangerous issue of false negatives, in which a controlled pathogen is misidentified as a benign sequence. We have observed this issue in practice as well, but do not report details here for reasons of informational safety.

Moreover, this situation is unstable, with the scope of the problem appearing to be expanding. Inspection of a few of the sequence dates from top matches found that most of the matched sequences, pathogenic or otherwise, were from the last few years. This is unsurprising, given that both the volume of sequencing data and the use of genetic tools are continuing to rapidly expand, and implies that degradation has been worsening and is likely to continue to do so even more rapidly. Worse yet, since the high numbers of tool sequences in the NCBI databases means there are often many sequences with identical or near-identical match scores, the actual set of accessions returned by BLAST may be volatile and change unpredictably.

The impact of BLAST degradation is particularly acute for the pragmatics in biosecurity and biosafety operations. The driving dynamics creating the issue mean that it is precisely the most commonly used tools that are likely to be misidentified as the pathogens of greatest interest, and vice versa. As a consequence, the cost of false positives and the risks from false negatives are amplified and are likely having a significant but unrecognized impact on the ongoing biosecurity and biosafety operations of industry, government agencies, and other organizations.

There are several potential paths for mitigation of this problem. First, the rules and curation procedures for taxonomic classification in NCBI and other affiliated databanks might be adjusted to better support pathogen classification, but this would conflict with a wide range of other usages for which these databanks have been designed. Specialized curation methods, such as the FLAN [22] and VADR [23] systems for viral pathogen curation, help with some aspects of data quality but cannot address the underlying interaction between BLAST and the categorization of chimeric material. One might also consider producing a variant of BLAST that is more suited for pathogen identification, e.g., by taking taxonomic weighting into account, but it is unclear whether this is actually feasible. Another approach is to switch to using existing databases that apply stringent taxonomic standards in curation, such as NCBI `refseq`, but doing so would drastically reduce coverage of variant sequences. Ultimately, however, it will likely be advisable to shift away from BLAST against NCBI and towards emerging methods that have been specifically tailored for pathogen identification, such as FAST-NA Scanner [24], ThreatSeq [25], SeqScreen [26], or SecureDNA [27].

No matter the potential path to mitigation, however, another important challenge illuminated by this study is the lack of a comprehensive standard and test set for evaluating the quality of pathogen identification. Current practice has largely defaulted to the use of BLAST against NCBI as a “gold standard” for pathogen identification, despite the fact that results from BLAST vary based on settings and are subject to many different possible interpretation strategies. Even if consensus could be reached on these issues, however, the results presented here demonstrate that continued use of BLAST against NCBI as a standard for identification is simply not sustainable. As a consequence, it will be important for stakeholders in pathogen identification to work together to develop a comprehensive standard for assessing the efficacy of pathogen identification systems.

Finally, although this work has focused specifically on pathogen identification, the underlying problems of taxonomic ambiguity and weighting are likely also affecting other applications that make use of NCBI databases. We therefore recommend that other NCBI user communities should perform their own case studies to determine whether there are similar issues in need of mitigation in their own applications.

## 4 Methods

Sequence analysis was run via the NCBI BLAST web interface (https://blast.ncbi.nlm.nih.gov/Blast.cgi) for protein (`blastp`) using the non-redundant protein sequence (nr) database and NCBI’s standard default parameter values. Taxonomic classification counts were determined using the BLAST taxonomy lineage report, counting all taxonomies whose top result tied for the maximum bit-score. Closest match was determined by maximum bit-score, with a sequence considered non-controlled if no controlled material scored higher than uncontrolled material (i.e., ties go to uncontrolled material).

A sequence in NCBI was considered to be controlled if its assigned taxa was a biological agent on either the Australia Group or US Commerce Control List. Sequences assigned to the polyphyletic taxa of “artificial sequences” (`NCBI:txid81077`) and “plasmids” (`NCBI:txid36549`) were not used for determination of control status, as these collections intentionally mix controlled and non-controlled materials.

## Supporting information

Supplementary S1 is a FASTA file containing the protein sequences used in this case study.

Supplementary S2 is a FASTA file containing protein sequences extended with random flanking sequences, used in the preparation of Figure 2.

Supplementary S3 is an Excel file containing the per-sequence results from the extension experiments.

## Supplementary information

Supplementary information is included with additional details on the study and its results:

- Supplementary S1 is a FASTA file containing the protein sequences used in this case study.
- Supplementary S2 is a FASTA file containing protein sequences extended with random flanking sequences, used in the preparation of Figure 2.
- Supplementary S3 is an Excel file containing the per-sequence results from the extension experiments.

## Acknowledgments

This document does not contain technology or technical data controlled under either U.S. International Traffic in Arms Regulation or U.S. Export Administration Regulations.

## Author Contributions

Conceptualization, Methodology: J.B., A.C., J.M. Investigation, Formal Analysis: J.B. Writing: original draft: J.B., review & editing: J.B., A.C., J.M.

## Footnotes

1 Throughout, we will use NCBI databases to refer to either NCBI’s databases or the equivalent ENA and DDBJ databases, which are maintained in synchrony with one another.

